# Species-specific sensorimotor noise levels explain saccadic suppression strength differences between macaques and zebrafish

**DOI:** 10.1101/2025.06.09.658731

**Authors:** Giulia Soto, Florian A. Dehmelt, Matthias P. Baumann, Ibrahim Tunç, Yue Yu, Tatiana Malevich, Ziad M. Hafed, Aristides B. Arrenberg

## Abstract

Active sensing necessarily requires integrating information about both self-generated movements as well as past, present, and future afferent inputs. However, such information is inherently variable and relies, at least in part, on uncertain extrapolations. Here, using saccadic suppression of visual sensitivity as a classic example of sensory-motor integration, we show that suppression strength in two drastically different species, macaque monkeys and zebrafish larvae, may be a direct outcome of efficient state estimation in the presence of uncontrollable sensory and motor variability. Bayesian estimator models suggest that optimal saccadic suppression should rely not just on the sensory-motor information being processed, but also on the time-dependent magnitude of unexplained variability in the nervous system encoding it. In both macaques and zebrafish larvae, and using matching visual stimulation regimes optimized for each species’ receptive field properties, we experimentally measured saccadic suppression strength in the superior colliculus (SC) of the monkeys and the homologous optic tectum (OT) of the fish. We also experimentally quantified additive and multiplicative motor noise components in the oculomotor behavior of both species, and we furthermore estimated noise in the sensory systems. We found that inter-species differences in sensory and motor noise levels and their theoretically predicted impacts on visual sensitivity are consistent with our experimentally observed differences in saccadic suppression strength between the fish and the monkeys. Because sensory and motor noise levels can reflect the amounts of available neural resources committed to a given task, our results strongly underscore the value of incorporating computational resource limits into investigating performance differences that have evolved in homologous brain areas.

## Introduction

When moving through the world, our brains constantly reconcile potentially conflicting information arising from sensory inputs in a changing environment and from the internal signals of our own body movements. One striking example of this non-stationary relationship between sensory and motor information is the active vision phenomenon of saccadic suppression^1,2^. Saccadic suppression refers to a brief reduction in visual sensitivity around the time of saccadic eye movements^3–9^ (Figure 1A). Saccadic suppression is a highly conserved phenomenon across a wide range of species, from zebrafish larvae^10^ to rodents^8,9^ and primates/humans^3,4,6,8,11,12^.

**Figure 1.**
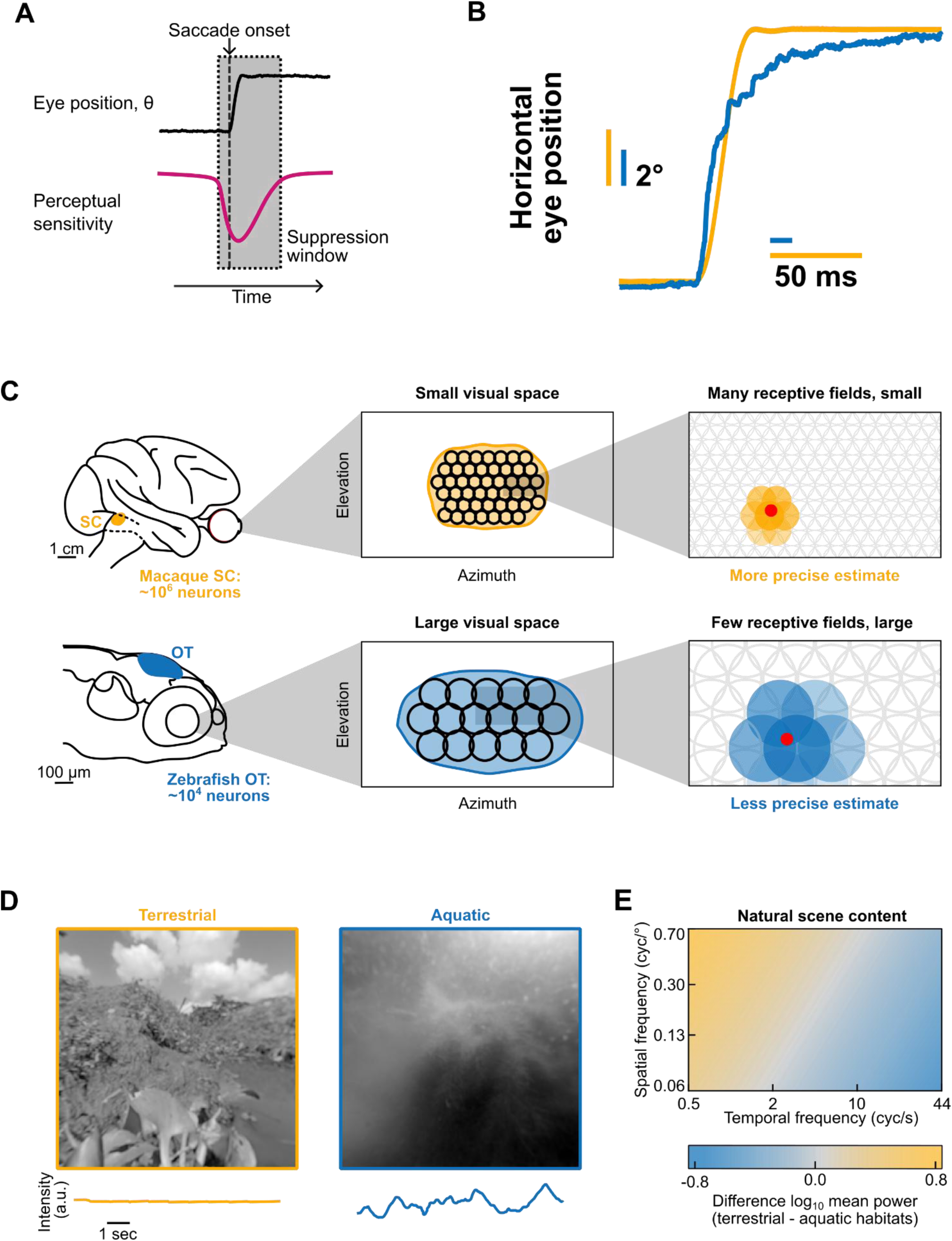
Saccadic suppression in the visual systems of two evolutionarily distant vertebrates, macaque monkeys and larval zebrafish. (A) In a suppression window around the time of a saccadic eye movement, visual perceptual sensitivity to exogenous stimuli is reduced. (B) Horizontal eye position during a saccadic eye movement of macaques (orange trace) and zebrafish (blue trace). The zebrafish saccade is executed more slowly and less accurately than the macaque saccade. (C) Schematic linking neuronal resources to accuracy of sensory estimation in macaque and zebrafish. Macaque superior colliculus (SC) contains ∼100 times as many neurons as the homologous zebrafish optic tectum (OT). This allows macaques to sample their significantly smaller visual field with many small-size receptive fields. In contrast, zebrafish have to fill a larger visual field with fewer, and larger receptive fields, leading to much coarser sampling of visual space. Thus, macaques can estimate the relative positions of visual stimuli (e.g. red dot) more precisely than zebrafish. (D) Example naturalistic images of terrestrial and aquatic habitats, encountered by macaques and zebrafish. Images were taken from Cai et al., 2023b^34^. Intensity traces below the images describe changes in luminance for each image (in arbitrary units) during a 10 sec time window. (E) Schematic depiction of difference in power spectra between terrestrial and aquatic habitats for spatial and temporal frequency content. Terrestrial scene content is dominated by high spatial and low temporal frequency content; aquatic scene content is dominated by low spatial and high temporal frequency content. Adapted from Cai et al., 2023a^35^.

Classically, saccadic suppression has been viewed as a perceptual problem, in which saccadic eye movements introduce large visual perturbations that the visual system needs to cope with. Recent work, however, has reframed saccadic suppression into the context of optimal sensorimotor control^13^. In this context, eye position control is conceptually modelled in a closed-loop control system, which dynamically weighs sensory and motor information, based on their inherent reliability, to optimally estimate the state of the eye. As a result, reduced sensory sensitivity is viewed a consequence of the need for instantaneous state estimation by the brain.

Four distinct sources of uncertainty are accounted for within such a Bayesian estimation framework: the sensory input, the dynamics of the eye plant, signal-dependent motor noise coming from the control signal itself, and the relay delay of sensory signals from the retina to higher computing areas^13^. Taking these variable sources of noise into account, specifically around the time of rapid eye movements, sensory signals are very unreliable in estimating the eye plant’s instantaneous state. This calls for a reduction in sensory gain, as can be observed during saccadic suppression, and it suggests that saccadic suppression is not merely a coping mechanism of the visual system, but in fact also a functionally beneficial component of overall motor control.

The above mechanistic explanation of saccadic suppression relies on anatomical structures that combine sensory and motor signals. In mammals, this is found in the superior colliculus (SC), and in lower vertebrates such as zebrafish, in the homologous optic tectum (OT)^14–18^. Studies in the macaque SC and zebrafish OT have shown that saccadic suppression can indeed be observed in neurons within these structures^6,10,16,19–21^, despite their large evolutionary distance from one another. This could be, at least in part, because brainstem circuitry underlying oculomotor control is relatively conserved between zebrafish and macaque^22–27^.

Despite such conservation, differences between zebrafish and primates allow us to consider the computational limits of saccadic suppression from an evolutionary optimization perspective. For example, due to a lack of cortex, and therefore the lack of important structures for saccade planning and generation, such as the frontal eye fields^28^, zebrafish saccades are in most contexts not goal-directed but rather reflexive (but see Ref.^29^). In zebrafish, the known descending control of saccades is limited to midbrain and diencephalic structures, such as the OT and the pretectum^30–33^. Further, the functional anatomy and natural scene statistics suggest that the zebrafish have only scarcer sensory information available (when compared to primates), which is aggravated by relatively poor eye position stability (Fig.1B, C, D, E). Thus, the smaller brain of zebrafish and the very different natural scene statistics provide us with an ideal opportunity to understand how computational limits in different brain sizes act to quantitatively reshape what are otherwise qualitatively identical phenomena in multiple species.

Starting from a theoretical foundation, we implemented a computational model allowing parameterization of different sensory and motor noise contributions to saccadic suppression strength. Then, using a matched visual stimulation approach (Methods), we experimentally measured saccadic suppression strength in homologous brain structures within macaque monkeys and zebrafish larvae, two species at opposite ends of the vertebrate evolutionary spectrum. Our results suggest a prominent role of sensory and motor noise in dictating saccadic suppression strength in the two species.

## Results

### Saccadic suppression as efficient sensory-motor estimation

Eye position control can be optimized in a closed-loop system, and a recent study suggested that saccadic suppression might be caused by efficient sensory-motor estimation across rapid eye movements^13^. According to this framework (Figure 2), a Bayesian estimator dynamically balances predictions from both sensory and motor information in an optimal way, according to the ever-changing reliability of such information, particularly across saccades. This framework can explain why time periods of visual uncertainty (i.e. during saccades) are associated with a (partial) suppression of the sensory representations in sensorimotor neurons. Anatomically, this state estimator can likely be found in the mammalian SC or the teleost OT, where neural correlates of saccadic suppression have been found^6,10,16^, and where spatial maps for oriented movement control have been documented^36–38^. While the two structures are anatomically homologous, neuron numbers - and therefore computational resources attributed to the respective anatomical structures in macaque monkey and zebrafish - vary greatly. Therefore, in what follows, we first estimated the implications of SC and OT neural resources in the two species on sensory and motor noise ranges, and we then investigated the impacts of such noise ranges on the experimentally observed and theoretically predicted strengths of saccadic suppression.

**Figure 2.**
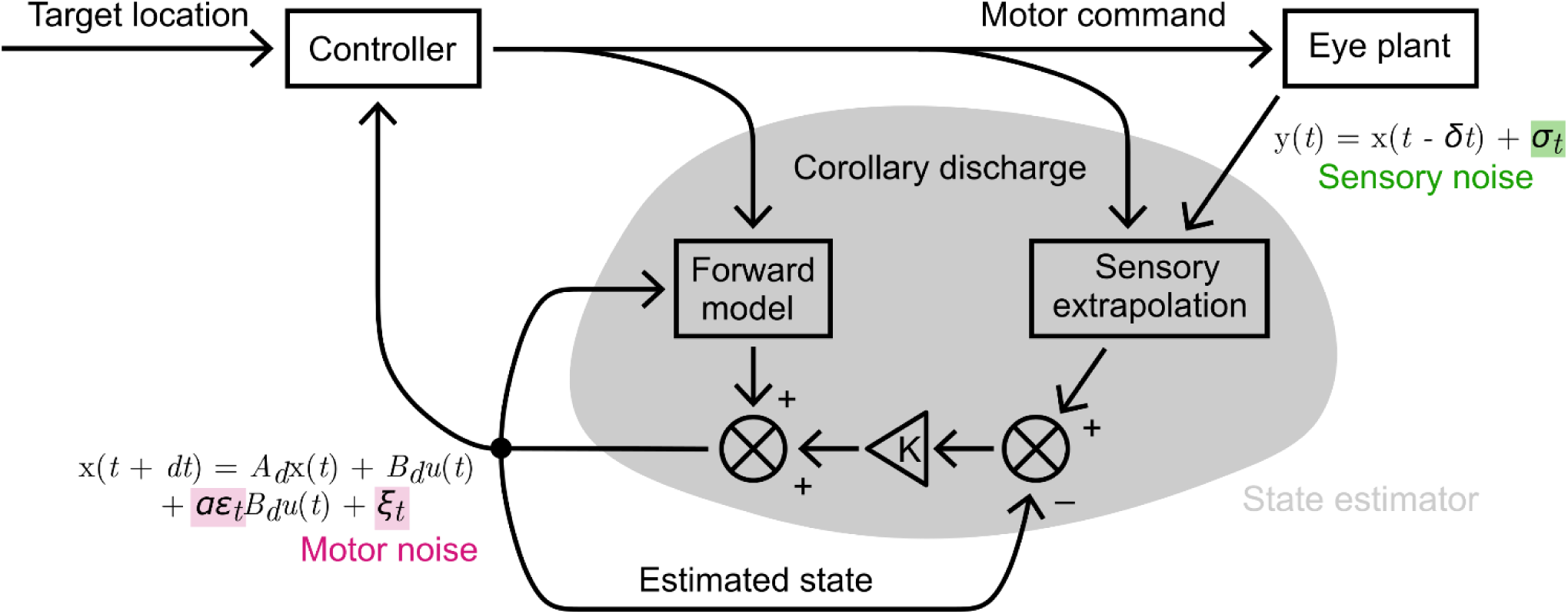
Saccadic suppression as efficient sensory-motor estimation. Schematic illustration of Bayesian state estimation for closed-loop control of eye movements (adapted from Crevecoeur and Kording^13^). To estimate the current position *x* of a visual target, its previous position is relayed as visual input *y* with delay *δt* and visual noise *σ_t_*, then combined with motor activity from the corollary discharge. This extrapolation with sensory information is weighted against a purely feedforward motor extrapolation via Kálmán filter *K*. Eye position is moved towards this estimated target by motor control signal *u* with multiplicative noise *ε_t_*, additive noise *ξ_t_*, and a scaling factor *α*. *A_t_* and *B_t_* capture the dynamics of the eye plant, *t* is time, *dt* the time step. Changes in *K* correspond to saccadic suppression or enhancement. See Methods for the model and notations.).

### Saccades across the species reflect different underlying neuronal resources

Despite the above-mentioned homologies between the SC and OT, the two structures are built with drastically different numbers of neurons. We estimated, based on brain volume and the total number of neurons in each organism, that the macaque SC has ∼100 times as many neurons as the zebrafish OT (Figure 1C, Methods). This difference in neuronal resources should be directly reflected in the oculomotor control of the respective species, as the two species markedly differ in their ability to execute precise saccadic eye movements. Consider, for example, Figures 1B, 3A, 3C: Eye position during saccades in zebrafish and macaques was not quantitatively identical, with zebrafish saccades being less accurate and having frequent over- and undershoots. Moreover, the eye movements had large variability in saccade offset times, indicating less precise motor control due to fewer neuronal resources. Further, when comparing sensory accuracy (Figure 1C), we noted that zebrafish have a much larger visual field compared to macaques, and with fewer neurons and larger receptive fields^39–42^. The visual field is thus only coarsely sampled in zebrafish, likely leading to less precise sensory estimates. In summary, the resource-constrained zebrafish oculomotor system, while anatomically conserved, may have to rely on much more uncertain visual and motor information than that of macaques, particularly peri-saccadically. As we show below, this has direct implications on the strength of saccadic suppression in the two species.

### Zebrafish motor and sensory signals are much noisier than those of macaques during saccadic eye movements

To assess the effect of noise levels on theoretical predictions of efficient sensory estimation, we first experimentally quantified motor noise before, during, and after saccades in macaque monkeys and zebrafish (Figure 3). We recorded saccade traces (see Methods, “Noise estimation”) in both species (Figures 3A and C) and estimated noise parameters for use in the model (Figures 3B, 3D). First, eye traces were split into pre-saccadic, saccadic, and post-saccadic sections (Figures 3B, 3D). Pre- and post-saccadic sections were used to determine additive motor noise, *ξ*, since it is modeled as independent of the motor control signal for saccades (see Methods, “Additive motor noise”). To quantify multiplicative motor noise, *ɛ*, we fit a sigmoidal curve to the saccadic section of the eye traces (Figures 3B and D). From the time derivative of the sigmoidal fit, we derived the control signal *u(t)*. A piecewise-constant approximation of *u(t)*, combined with the previously determined additive motor noise, *ξ*, was then used to estimate *ε* (see Methods, “Multiplicative motor noise”). We thus determined that zebrafish motor control is much noisier than that of macaque monkeys, with the additive motor noise being about 10 times higher in zebrafish, and the multiplicative motor noise being nearly 40 times higher (Figure 3E). In our estimation of motor noise, we excluded saccades with significant over- or undershoots from our analysis (see Methods – Noise estimation), since they are not the object of this study. Importantly, though, relative motor noise estimates between species are not dependent on this exclusion of trials and persist when including all saccade types.

**Figure 3.**
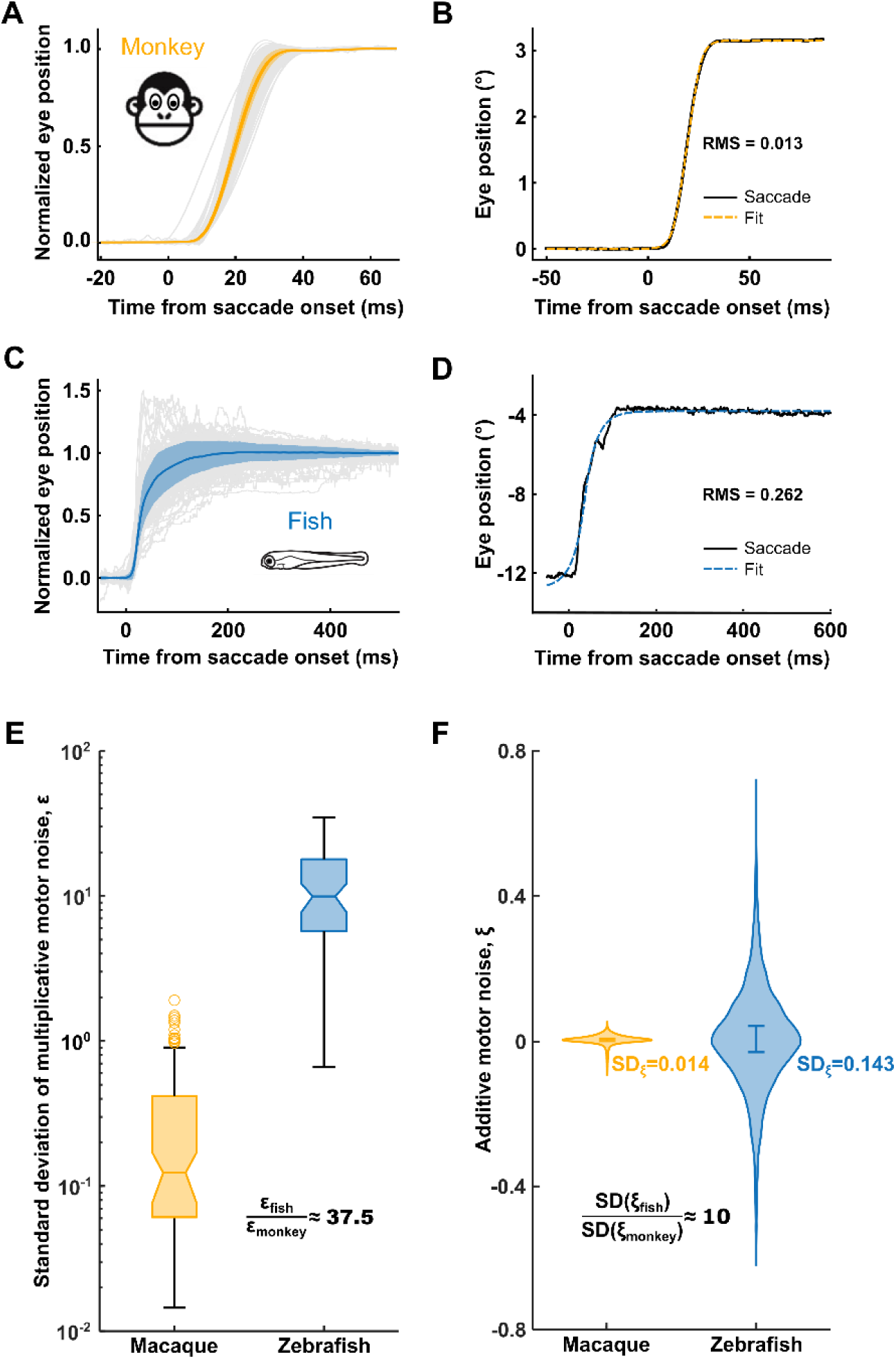
Zebrafish motor signals are much noisier than those of macaques during saccadic eye movements. (A) Normalized eye position during saccadic eye movements in macaques. Grey lines indicate individual trials, colored line indicates average eye position and shaded envelope indicates standard deviation. Saccades were normalized to saccade amplitude, determined by the difference between saccade onset and offset position (Methods). *N = 1,445 saccades* (B) Example macaque saccade with sigmoidal fit (dashed line). Deviations from the fit (RMS) are considered motor noise (for detailed description see STAR Methods). (C and D) Same as (A) and (B) for zebrafish. The eye movements were much noisier than for macaques. *N = 208 saccades* (E) Standard deviations of estimated multiplicative (ε) and additive (ξ) motor noise. Boxplot shows standard deviations of multiplicative motor noise estimates for multiple velocity bins for macaque and zebrafish during saccadic eye movement (see Methods). Central line indicates median, box encloses 25th to 75th percentile and whiskers extend to minimum and maximum of data; circles show outliers. (F) Violin plot shows distribution of additive motor noise estimates pooled across all pre- and post-saccadic stationary phases for macaque and zebrafish; error bars indicate the standard deviation.

While the motor noise was easy to quantify experimentally, estimating sensory noise is more challenging. This is because it is not known which visual neurons macaque or zebrafish use to estimate their own eye positions. Therefore, instead of estimating sensory noise using computational models of image-based eye position inference - which would require assumptions regarding the identity of the relevant neuronal population in each species - we decided to estimate sensory noise levels based on three mechanistic possibilities that each point towards sensory noise levels being about 10x larger in zebrafish compared to macaques. In the first mechanistic scenario, the precision of sensory information increases with the number of available neurons. Visual receptive fields oftentimes tile visual space evenly (as illustrated in Figure 1C), meaning that different regions of visual space activate similar numbers of neurons (we ignore the fovea and the area temporo-ventralis/area centralis of macaques and zebrafish here). The relation of the sensory noise parameter *σ* for both species can then be estimated from the number of neurons in their respective visual brain areas (see Methods, “Estimation of neuron count”, “Sensory noise”). There are about 1 million neurons in the SC^43,44^ and 10,000 neurons in OT^41,42^. This ratio of 100 translates to a factor of 10x, when taking into account two-dimensional tiling of the visual field with receptive fields.

For the second scenario that is closely related mathematically, consider that eye position is inferred using a winner-takes-it-all mechanism, in which only the best-activated neuron conveys the information about the position of an object in space relative to the eye^44^. In this scenario, eye position estimation is as precise as the receptive field size of individual neurons. While zebrafish receptive fields in the OT range from approximately 5° to 130°^45–48^, macaque receptive fields in the SC range from 0.5° to 20° according to Chen et al. 2019^49^. Therefore, also in this scenario, sensory noise should be approximately one order of magnitude higher in the zebrafish visual system than for macaques.

Finally, in the third scenario, we acknowledge that visual inference of eye position is only possible, when a structured visual environment can be seen by the animal. Previous studies on natural stimulus statistics^35^ have shown that aquatic environments contain fewer structures of high spatial frequency than terrestrial habitats. Furthermore, there is more (temporal) luminance noise in aquatic environments due to sunlight refraction at caustic water surfaces and due to floating particles in the water, both leading to higher temporal frequencies of visual stimuli in the aquatic environment (Figure 1D, 1E). Cai et al. showed that spatial frequencies between 0.1 and 0.7 cycles per degree were about 10 times more abundant in terrestrial environments, and temporal frequencies in the range of 5 to 44 cycles per second were about 10 times more abundant in aquatic environments (cf. Fig. 4G in Cai et al., 2023a^35^). This temporal noise and these spatially less-contoured natural stimuli in the aquatic environment should impede precise eye position inference via the visual system. On the other hand, the contrast sensitivity function of macaques reaches much higher spatial frequencies^50,51^, again pointing to lower sensory noise in these animals.

**Figure 4.**
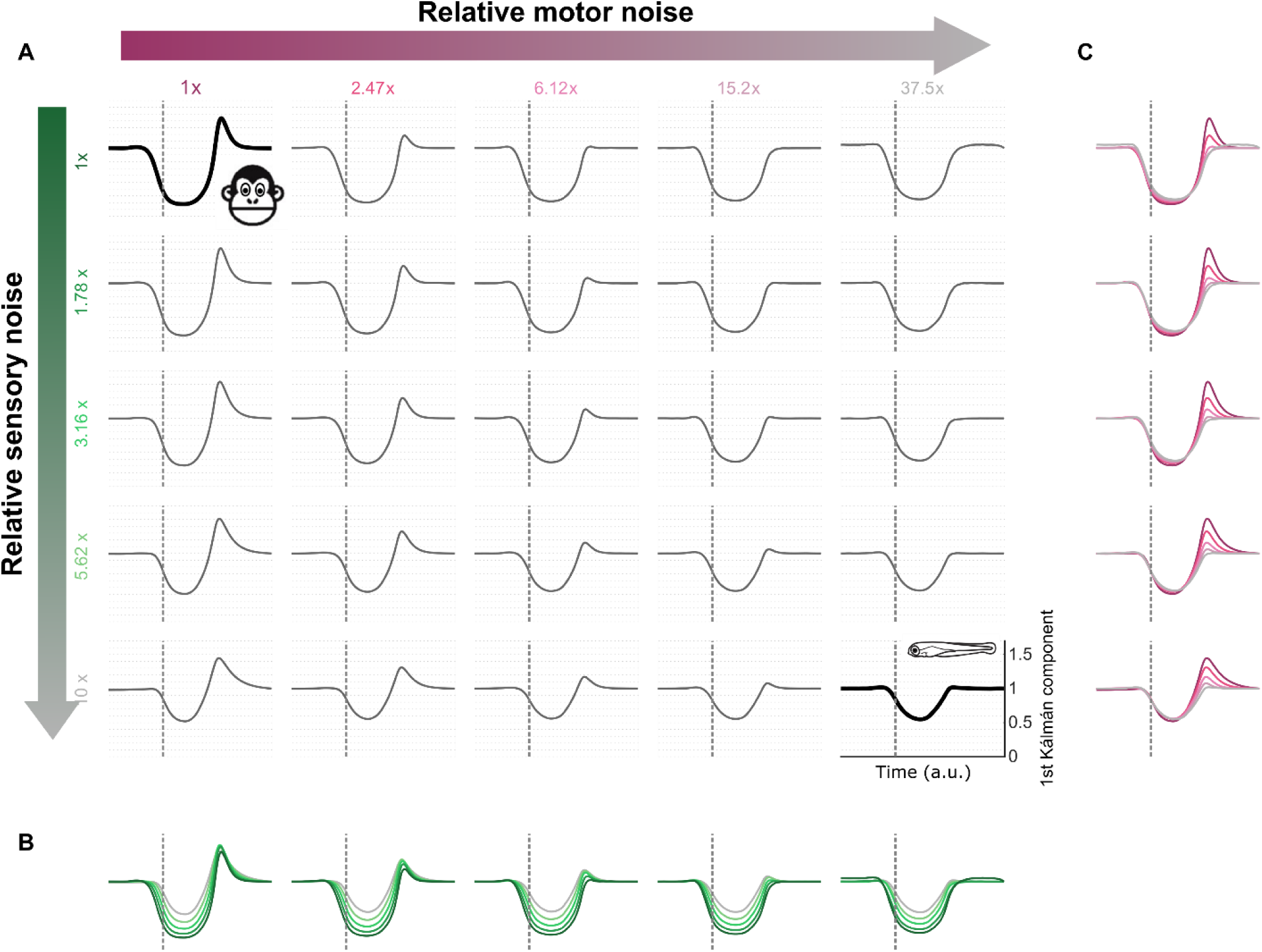
Bayesian estimation model predicts differences in saccadic suppression profiles across the range of noise parameters observed in macaque and zebrafish. (A) Optimal time course of saccadic suppression predicted by the model. All solid black traces show the first element of the optimal Kálmán matrix; vertical dashed lines show saccade onset. Different combinations of sensory and motor noise entail different amounts of saccadic suppression. The thick trace on the upper left was obtained for macaque-like noise parameters, and the thick trace on the bottom right for zebrafish-like values. All noise parameters shown here (e.g., “10x”) are relative to macaque values (“1x”). Species differences in saccade length can be taken into account through corrective terms, slightly lowering the zebrafish-like parameters (see Methods). (B) Increased sensory noise reduces the depth of saccadic suppression across motor noise regimes, but in the parameter range investigated, hardly affects its overall shape. Curves are superpositions of those in the columns of (A), with shades of green indicating different magnitudes of sensory noise for the same motor noise. (C) Increased motor noise discourages post-saccadic enhancement, and this effect is robust across sensory noise regimes. Curves are superpositions of those in the rows of (A), with shades of magenta indicating different magnitudes of motor noise for the same sensory noise.

Based on the above-described three mechanistic possibilities, we estimated that the sensory noise for sensorimotor estimation might be ten times noisier in zebrafish compared to macaques. As we show further below, we also simulated other ranges of sensory noise in the model, allowing us to more strongly support our conclusions regarding inter-species’ saccadic suppression strength levels.

After measuring the additive and multiplicate motor noise levels experimentally, and estimating the inter-species sensory noise difference, we were now in a position to generate predictions on saccadic suppression strength across the two species.

### Suppression strength differences across species are consistent with their noise levels

We first identified model parameters to reproduce the curve predicted in the original study^13^. We decided to covary additive and multiplicative motor noise in our model; that is, we made *ε* = *ξ* (Methods). This was justified due to the high level of abstraction of this model, where sensory and motor information are modeled as abstract variables without reference to specific neural implementations. It was also further justified because across species, the ratios of species’ noise levels were on similar orders of magnitude for both additive (*ξ*) and multiplicative (*ε)* motor noise (Figures 3E and 3F). Then, we ran further simulations in which we tested the effects of varying sensory and/or motor noise by factors of up to 37.5 times, reflecting the inter-species noise ratios inferred from our experiments (Figures 1C, 3E, 3F, 4A). For simplicity, we did not explicitly represent saccade duration differences between species, but this can be done and would slightly reduce these factors without affecting the conceptual outcome (see Methods, “Noise estimation”).

Higher sensory noise indeed reduced the predicted optimum of saccadic suppression strength, consistent with the notion that if sensory information is always unreliable, then saccades have less effect on the trade-off involved in efficient state estimation (Figure 4B). The effect of motor noise, on the other hand, was not symmetric to that of sensory noise, and noise levels observed in zebrafish primarily affected predictions of post-saccadic enhancement (Figure 4C; see below and Fig. 5 for what we mean by post-saccadic enhancement). This result is again congruent with our intuition when comparing post-saccadic stability of eye position across the two species. As visual uncertainty in larval zebrafish is still high, even after the saccade has completed, corrections based on the short-term enhancement of post-saccadic visual information are less effective in zebrafish. Macaques, with much more stable post-saccadic eye position (Fig. 1B), however, benefit from a brief post-saccadic enhancement. In fact, snapshot enhancement, or improved feature estimation, for a brief moment immediately following a saccade, has been shown in multiple biological systems^52–54^. Thus, if macaque monkeys have lower sensory and motor noise than zebrafish, then the results of Figure 4 predict much stronger saccadic suppression, as well as post-saccadic enhancement, in the monkeys than in the fish.

**Figure 5.**
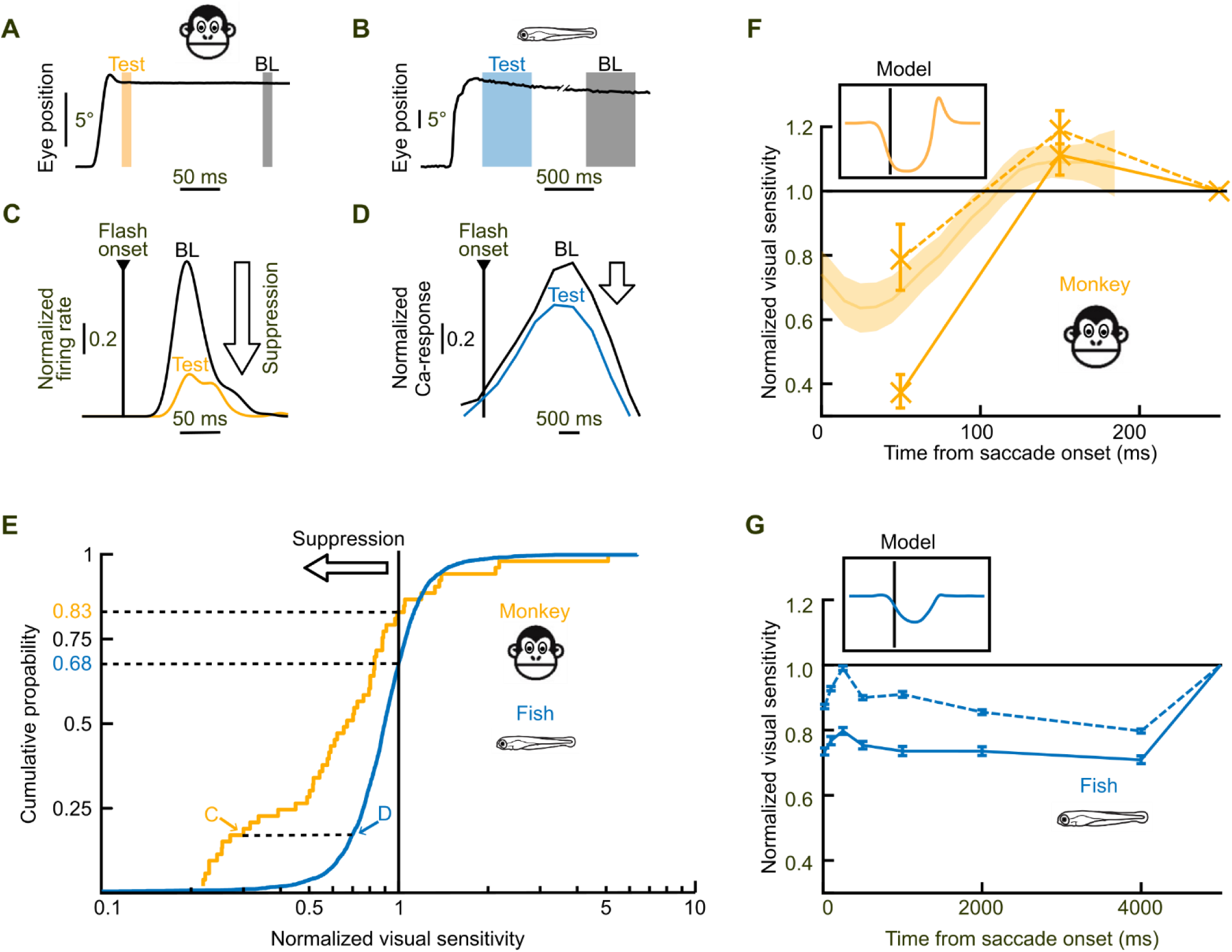
Saccadic suppression in the monkey superior colliculus and the zebrafish optic tectum matches model predictions. (A) Example eye trace from a monkey with the timing of the baseline flash (“BL”; ∼200 ms after the saccade) and a test flash (“Test”) ∼50 ms after saccade onset. (B) Example eye movement trace from a zebrafish, illustrating baseline and test flash timing. Note: only a test flash 250 ms after saccade onset is shown here; in panels (E) and (G), test flashes at 20, 100, and 250 ms were pooled. (C) Example neuron from monkey SC: visual response to the baseline flash (black) and to the test flash (orange). There was strong saccadic suppression. (D) Example neuron from zebrafish tectum: visual response to baseline flash (black) and pooled test flashes (blue). Saccadic suppression was present, but weaker than in the example monkey SC neuron. (E) Cumulative distribution functions of suppression strength across neurons. Example neurons from (C) and (D) are highlighted. 83% of SC neurons (orange) show suppression at 50 ms post-saccade; 68% of OT neurons (blue) show suppression across the delays 20, 100 and 250 ms. Average suppression is stronger in SC neurons, than in tectal neurons (p < 0.001, Wilcoxon rank-sum test). (F) Time course of suppression in monkey SC. The faint continuous curve indicates data from Bellet et al. 2017^55^ in which a full time course was measured (with microsaccades); crosses show the current data from all neurons (dashed line) and from the significantly suppressed ones (solid line). The dashed line closely matches the Bellet et al. 2017^55^ time course, which also included all measured neurons. Importantly, we saw systematic post-saccadic enhancement, directly consistent with the model predictions. Inset: model prediction for low sensory and motor noise, as in Figure 4A, top left. (G) Suppression time course for zebrafish OT neurons, analogous to (F). Inset: model prediction for high sensory and motor noise, as in Figure 3A, bottom right.

To test for such cross-species differences in saccadic suppression strength, we recorded neural activity in the SC of macaque monkeys and the OT of zebrafish. We matched the visual stimulation parameters between the two species as closely as we could under some technical constraints. Specifically, for both species, we showed visual probe flashes on a textured background and measure neuronal responses to said probe flashes within the SC or OT. Critically, the textured background statistics were optimized for each species’ visual acuity resources (Methods): spatial frequency ranges were matched to the receptive field sizes expected in the respective animal, which is a similar approach to our earlier multi-species comparison of saccadic suppression^8,9^. SC activity was measured using electrophysiological recordings of linear multi-electrode arrays; OT activity was measured using 2-photon calcium imaging. Then, we compared probe flash responses shown in temporal isolation (referred to as “Baseline flash” in Figures 5A-D; Methods) to responses of probe flashes shown peri-saccadically (“Test flashes” in Figures 5A-D). With saccade duration in macaques being 24-45 ms (mean = 37.6 ms) and saccadic suppression lasting approximately 40-60 ms^6,8,16^, we chose to show our baseline flash at 200 ms after saccade onset for macaques to ensure no signal interference from the saccade. For zebrafish, suppression is known to last much longer^10^ (4 s), which is why we showed the baseline flash at 8 s after saccade onset. Probe flashes for zebrafish were longer to allow for signal integration of the relatively slow calcium indicator. Additionally, since calcium imaging recorded multiple neurons at the same time and had a worse temporal resolution than electrophysiology approaches, we used a whole-field flashes for zebrafish and local flashes covering the receptive fields for macaques.

Figures 5A and B schematically illustrate the timing of visual stimulus presentation relative to saccade onset in both species. Example neurons from the SC and OT are shown in Figures 5C and D, respectively. In both cases, responses to the flash were markedly reduced when the stimulus appeared shortly after a saccade, indicating clear saccadic suppression at the level of single units in both primates and fish. To compare suppression strength across the population, we computed cumulative distribution functions (CDFs) of suppression indices (Methods) for all included neurons (Figure 5E). The distributions revealed a clear species difference: saccadic suppression in the macaque SC was generally significantly stronger (statistical results are detailed in the legend of Figure 5) and more prevalent than in the zebrafish OT, consistent with the model predictions (Figure 4) that higher motor and sensory noise in zebrafish reduce the benefit of discounting sensory inputs during and immediately after saccades. Thus, the existence of a difference in suppression strength for zebrafish^10^ and macaque monkeys^6,16^ can be explained by an optimization principle related to efficient state estimation under the presence of noise.

Finally, in both species, we experimentally tested visual sensitivity at multiple times relative to saccades (Methods). This allowed us to further compare the empirical data to the predictions of our model, and especially with respect to post-saccadic enhancement of visual sensitivity (Figures 5F and 5G). For both species, the model reproduced the fact that there is always an extended time period in which visual sensitivity is reduced around saccades, regardless of suppression strength. Importantly, besides correctly predicting that saccadic suppression would be stronger in monkeys than in fish – driven solely by species-specific noise levels estimated independently (cf. Figure 3) – the model also predicted that post-saccadic enhancement (Fig. 5F and Bellet et al., 2017^55^) is more likely in monkeys than in fish. As mentioned above, this reflects the markedly different post-saccadic stability of eye position in the two species (Fig. 1B).

In summary, we were able to experimentally reproduce both predictions that the computational model made regarding the effects of sensory and motor noise on efficient sensory-motor estimation during saccades. First, increased sensory noise in zebrafish led to weaker saccadic suppression strength. Second, increased motor noise led to reduced post-saccadic enhancement. In our view, the alignment between modeling and data in our study suggests that efficient sensory-motor estimation under species-specific noise constraints provides a unifying account of saccadic suppression strength across vertebrates.

## Discussion

Saccadic suppression is a highly conserved active vision phenomenon, that has been observed across multiple species^8,10^. Here, by combining computational modelling with neurophysiological recordings in homologous structures, using matched visual stimulation regimes in two evolutionary distant species, macaque monkeys and larval zebrafish, we showed that saccadic suppression can be understood as optimal state estimation under species-specific noise constraints. Specifically, a Bayesian state estimation model predicted weaker peri-saccadic suppression with increased sensory noise and weaker post-saccadic enhancement with increased motor noise. Species-specific noise estimates and neurophysiological recordings in macaques and zebrafish confirmed these predictions biologically. Together, these observations support the view that saccadic suppression is not merely a visual coping mechanism, but rather an adaptive strategy for state estimation under resource constraints.

One implication of our findings is that saccadic suppression may be better understood not as an exclusively visual phenomenon, but rather in a broader context of sensorimotor optimizations that expand beyond the visual domain. For example, sensory gating of the tactile system in mouse whiskers, where responses to exogenous stimuli are reduced during self-motion, have been described^21,56,57^. Similarly, in the tactile system of cats^58^, humans^59^, and non-human primates^60,61^ somatic sensory gating during voluntary movement has been found. Sensory gating further expands to the proprioceptive system in primates^62^, as well as the orofacial^63^ and the auditory system in humans^64^. Together, these observations suggest that a transient reduction in sensory gain during self-generated action (such as touch, arm/body movement, or speech) may represent a general computational strategy for efficient sensorimotor control.

While the existence of sensory gating seems to be broadly conserved across sensory modalities and species, the strength of such gating is likely dependent on species-specific neuronal resource constraints and ecological demands. In the case of saccadic suppression, macaques rely heavily on foveated vision with dense receptive field sampling and precise goal-directed saccades for visual perception. Under these conditions, brief sensory reduction during periods of elevated sensorimotor noise, followed by brief sensory enhancement after the completion of a precisely executed saccade, may bring substantial benefits for efficient sensory estimation. Zebrafish, on the other hand, have a much broader visual field, which is less densely sampled, and oculomotor control is significantly less precise, compared to macaques. Consequently, uncertainty in the oculomotor system of zebrafish is high, peri-saccadically or else. Therefore, there is little benefit to reducing sensory input during saccades or enhancing post-saccadic snapshots. Additionally, natural aquatic environments (in comparison to terrestrial ones) contain fewer high spatial frequency features and more temporal luminance fluctuations^35,65^, further contributing to the relative uncertainty in the zebrafish oculomotor system. Altogether, our findings suggest that quantitative differences in saccadic suppression reflect adaptations to species-specific neural resource constrains as well as adaptations to the statistical structure of the environment, the animal evolved in.

While the Bayesian state estimation model captures species-specific differences in suppression strength well, it is agnostic to suppression length, as it operates in arbitrary time units. Indeed, suppression duration in macaque SC lasted less than 100 ms, whereas suppression in zebrafish OT persisted considerably longer (4 s). Previous work demonstrated that this prolonged suppression in zebrafish is not an artifact of experimental design but rather the effect of stimulus saliency, with behaviorally salient stimuli exhibiting considerably shorter lasting saccadic suppression^10^. One potential explanation for the inter-species difference of suppression duration could be the lower saccade frequency in zebrafish, compared to macaques. If saccades serve as a means to acquire novel information in the form of “snapshots”, then the temporal profile of sensory suppression should scale with the inter-saccade interval (∼180 spontaneous saccades/min in free-viewing macaques^66^, ∼2 spontaneous saccades/min for larval zebrafish in this study). Thus, animals that sample their environment more frequently, benefit from shorter lasting suppression, whereas animals with low sampling rates may need longer lasting suppression. Future work will have to determine whether saccade frequency shapes the temporal profile of saccadic suppression.

### Limitations of the study

Several limitations must be considered when interpreting the present findings. First, direct neurophysiological comparisons between macaques and zebrafish are inherently challenging, due to the necessary differences in technical experimental design across species. Neural activity was measured using electrophysiology in macaques and calcium imaging in zebrafish, which lead to differing temporal resolution. Visual stimulation paradigms had to be matched under consideration of the visual field properties of each animal. For macaques, with forward facing vision, we used a simple screen and used local probe flashes, directly matching the receptive field of the neuron of interest. In zebrafish, which have a much larger visual field, we used a full surround visual stimulation arena and global stimuli, since multiple neurons were recorded in parallel and the suboptimal temporal resolution of calcium imaging prevented us from first measuring the receptive field of individual neurons. Soto et al. 2025^10^ show that the strength of saccadic suppression is even smaller in zebrafish, when a local stimulus (moving dot) was used, which suggests that the species’ difference in suppression strength might even be larger than shown here. Furthermore, saccades were visually guided in macaques and spontaneous in zebrafish. Saccade size also varied greatly between the two species, with zebrafish routinely performing saccades >20°, while the guided saccades in macaques were 7.5° large. While there is some evidence that saccadic suppression strength scales with saccades size in zebrafish^10^, the effect is small and not observed in macaques (Fig. 5F shows similar SC suppression for microsaccades and larger saccades) and thus does not affect the results of this study. While these methodological differences may have influenced the absolute magnitude and temporal profile of suppression measured, the central claim of weaker suppression and weaker post-saccadic enhancement in zebrafish was likely not affected.

The second limitation is regarding the sensory noise estimation. While motor noise could be directly inferred from saccadic eye traces, sensory noise could only be estimated indirectly from neuron numbers, receptive field properties, and/or naturalistic image statistics. However, since we have multiple lines of argumentation all converging on a relative difference of sensory noise on the scale of one order of magnitude, we are confident that the conceptual predictions of our model and our conclusion resting on it do not depend on the exact parameter choice. Also, the presented effects of noise in the model do not depend on the precise level of inter-species noise ratios chosen (cf. diagonal in Figure 4). Also, we did not model the experimentally known differences in saccade duration either explicitly or by correcting for different simulation time steps, potentially overestimating individual noise samples by a factor of two or more (see Methods, “Estimation of noise parameters”). This simplification did not affect our results conceptually. Further studies are needed to investigate the temporal aspect of saccadic suppression across species, i.e. whether longer durations of saccadic suppression and saccades (as observed in zebrafish) can be understood as adaptations to higher levels of noise.

And lastly, the Bayesian state estimation model employed here operates on a high level of abstraction and does not capture the biological implementation of saccadic suppression. Therefore, it cannot explain any species-specific circuit mechanisms, suppression duration, or the neuronal populations involved. Instead, the model should best be viewed as a conceptual framework, linking sensory and motor uncertainty to sensory gain modulation. Regardless of the high level of abstraction, the model still successfully captured the conceptual differences in saccadic suppression between two evolutionary distant species. We suggest that theoretical considerations of the implications of computational resource limits, like we did here, will also be valuable for our understanding of a wide range of further multi-modal, integrative brain processes.

## Methods

### Animal Care

All non-human primate and zebrafish experiments were evaluated by ethics committees and approved by the Tübingen regional government. For macaques we recorded SC activity from two adult, male rhesus macaque monkeys (A and F), aged 13 to 14 y and weighing 9 to 13 kg. For zebrafish we recorded calcium activity of 7 larval zebrafish 5-7 days post fertilization (dpf) with pan-neuronal GCaMP6f expression and homozygous for the *mitfa* mutation (*Tg(HuC:H2B-GCaMP6f)jf7; nacre -/-*)^67^. For eye position recordings we used 12 *nacre -/-* larvae, 5-6 dpf. Larvae were raised in E3 solution at 28°C on a 14/10 light/dark cycle.

### Noise Estimation

#### Macaque eye position recordings

The eye data used for the noise estimation is a subset of the data published in Hafed et al., 2019^68^. It comes from the delayed saccade task, in which the monkey fixated a central spot while a peripheral target (1-degree-diameter circle) appeared and remained visible. After a variable delay (500–1,000 ms), the fixation spot disappeared, cueing a saccade to the target. We used 1,445 trials involving four oblique saccade directions with varying amplitudes. Eye data acquisition and saccade detection were performed similarly to the procedures described below for the current monkeys’ task.

#### Zebrafish eye position recordings

Eye position of larval zebrafish was recorded at 1 kHz with a high-speed camera (iN8-S1 IDT, Integrated Design Tools, Inc., Pasadena, CA, USA) mounted on a stereomicroscope. Spontaneous saccades were detected using a motion trigger and 1000 frames recorded, including 200 frames pre-trigger. Recordings were manually screened for false triggers caused by non-saccadic motion within the frame and, if necessary, discarded. Eye position was extracted post-hoc using the ZebEyeTrack virtual machine^69^.

#### Additive motor noise

The zebrafish dataset was further filtered using strict quality control criteria because zebrafish saccades were spontaneous, unlike the target-driven saccades in the macaque dataset. To ensure only robust and voluntary saccadic movements were analyzed, zebrafish trials meeting any of the following criteria were excluded prior to analysis: saccades were not conjugate (57 trials excluded), saccade amplitudes were less than 5° for both eyes (67 trial excluded), and traces showed strong fluctuations for either eye, defined as a saccade having a local maximum with a prominence greater than 30% of the total saccade amplitude (26 trials excluded). The remaining traces were visually inspected, and 5 further traces clearly showing no saccade were manually discarded, leading to a total of 209 saccade traces.

In each trial, saccade onset and offset were detected with a velocity criterion matching the variability of the eye trace^70^, and trials split into pre-saccadic, saccadic, and post-saccadic periods accordingly. Additive motor noise was estimated from pre- and post-saccadic data only by fitting one affine function each to the pre- and post-saccadic traces of each trial using the Levenberg-Marquardt algorithm to minimize the root mean squared (RMS) error. The resulting RMS error, representing the standard deviation of the fit residuals, was used to estimate *ξ*.

#### Multiplicative motor noise

Multiplicative motor noise is associated with the control signal and thus estimated from saccadic periods under the assumption that the (previously estimated) additive motor noise remained stationary. We fit a sigmoid function to the saccadic part of each eye trace using a soft ramp model^71^ with the same fit procedure described above. As larvae occasionally showed pre-saccadic drift as well as large saccadic overshoots or undershoots, and these were not the object of the present study, such trials were excluded on a data-driven manner. Saccades with a fit score less than the average fit error minus 0.5 times the standard deviation of the fit error were considered, yielding ∼31% (64 out of 209 saccades) of the zebrafish dataset. For comparability, macaque saccades were also discarded via the same exclusion criterion, leading to ∼33% (474 out of 1445 saccades) of the dataset being used for the analysis. The exclusion threshold was high enough to ensure that both high-noise and low-noise trials with sigmoidal saccades were included, and only significantly distorted saccades excluded. For each trial, we then computed the time derivative of the best fit to estimate control signal *u*, which was pooled over all eye traces. To facilitate estimation of *ε*, we converted *u* into a piecewise-constant function by dividing saccades into time bins of 100 values, computing an average ū per bin, and discretizing it. Across trials, eye positions with the same discretized ū were pooled; the standard deviation of their distance from best fits provided one estimate of multiplicative noise per value of ū. As we observed no characteristic differences between estimates obtained under high or low control signals, we averaged them to obtain *ε*. The arbitrary scaling parameter *α* was kept at 0.06 throughout, as this allowed us to reproduce the original model results^13^.

#### Simulation time steps

In return for numerical simplicity, we did not explicitly model saccade duration. As a side effect, this imposed identical time steps in arbitrary time units on all simulations. This risks exaggerating parameter differences and, if quantitative temporal precision is required, should be accounted for by correcting noise samples by the square root of the step size in real-world time units, as for any other Wiener process. As it stands, zebrafish-like parameters may be overestimated by at most the square root of the ratio between saccade durations. In the case of a 20 ms saccade in macaque vs. 100 ms in zebrafish (Fig. 3A, C), we would need to reduce noise samples by a factor of 2.24; in such a case, the predicted suppression for zebrafish would more closely resemble that for the parameter pair 5.62x/15.2x shown in Figure 4A than the one currently highlighted, but without affecting the result conceptually.

#### Normalization of eye positions

Saccade traces in Figure 2 were normalized to an average range of 0 to 1. First, start and end eye positions were calculated as the average positions during the 50 ms windows immediately preceding saccade onset and following saccade offset, respectively. The start position was then subtracted from each trace, and the result was divided by the saccade amplitude, defined as the difference between the end and start positions.

#### Estimation of neuron count within brain areas

Our estimation of sensory noise depended on the difference in number of neurons in primate superior colliculus versus zebrafish optic tectum. The SC of macaques has an estimated volume of 15 mm^3^ (cf. Figure 3D in Chen et al., 2019^49^, we used the raw data of the SC surface reconstruction). The average macaque brain has a volume of 90.000 −100.000 mm^3^ and an estimated number of 6 x 10^9^ neurons^43,44^. Assuming the neuronal density to be roughly homogeneous, we multiplied the total neuron number by the fraction of SC volume relative to whole brain volume, roughly totaling 10^6^ neurons. Note, however, that the homogeneous density assumption is a simplification, since subcortical structures have been reported to have lower neuron density than cortical structures^44^. 7-day-old zebrafish contain about 116,000 neurons in their entire nervous system^41^; tectal volume makes up about 8% of total nervous system volume^42^. Assuming, again, neural density to be roughly homogenous, we estimated that the tectum contains roughly 10,000 neurons. Based on our estimated neuron count and assuming, for the sake of argument, a uniform two-dimensional tiling of visual space, the 100-fold difference in neuron count translates into a difference of visual resolution on the order of *σ* = √100 = 10.

#### Sensory Estimation Model

To model the effect of noise on efficient sensory estimation, we reconstructed a closed-loop control model that had previously been used to explain the existence and time course of saccadic suppression^13^. This abstract model treats sensory and motor information as simple 4D variables (position and velocity of eye and target, respectively), without any reference to specific neuronal implementations. In addition to our own, the original authors kindly provided unpublished MATLAB code. We manually identified parameters to reproduce the originally published time course and to improve the stability of simulations across a wider range of noise levels. Final parameters (all unitless as per the model) included *ξ* ∊ [0.01, 0.375], σ ∊ [10^−6^, 10^−5^], scaling factor *α* = 0.06, sensory delay = 0.1, initial fixation time = 0.5, movement time = 0.05, stabilization time = 0.25, cost function tolerance = 0.01, initial target position = −10, final target position = 20, discrete time step = 0.02. Because inter-species differences were on a comparable order of magnitude in both *ξ* (37.5x) and *ε* (10x), and because the effect of multiplicative motor noise *ε* was relatively minor, we covaried the two types of motor noise in all simulations reported here, with *ε* = *ξ*. Because of the model’s high degree of abstraction, we cannot directly substitute our experimentally determined noise values. Therefore, we chose an approach, where we first reproduced the curve from the original modeling study^13^ and then multiplied the used noise levels to investigate the effects on the saccadic suppression curve (Figure 3).

### Macaque Electrophysiology

#### Experimental setup

The experimental setup closely followed the procedures outlined in our previous study^72^. In brief, experiments were conducted in a darkened room with the animal positioned approximately 72 cm away from a calibrated and gamma-linearized CRT monitor (covering ∼30° horizontally and ∼23° vertically). Stimulus presentation and data collection were managed using a custom-modified version of PLDAPS^73^, integrated with the Psychophysics Toolbox^74^ and interfaced with an OmniPlex data acquisition system (Plexon, Inc.).

Animal preparation for behavioral training and electrophysiological recording was carried out as described in earlier work^72^. Briefly, each animal was surgically implanted with a head post for head stabilization and a scleral search coil in one eye^75^ to enable high-precision eye tracking via electromagnetic induction^76^. A recording chamber was also implanted along the midline, tilted 38° posterior to vertical to target the superior colliculus. We used linear multi-electrode arrays (V-Probes, Plexon Inc.) with either 20 or 24 channels and 50 μm inter-electrode spacing to record neuronal activity. We minimized sampling bias by offline spike sorting.

#### Behavioral task

In each trial, monkeys performed a visually guided saccade task against a background of either a fine or a coarse texture. For each texture condition, one of 80 pre-generated images (80 fine, 80 coarse) was randomly selected and displayed as the background.

Each trial began with the presentation of a fixation spot, located 7.56° to the left or right of the screen center. After the monkey acquired and maintained fixation on this spot, it disappeared and simultaneously reappeared at the center of the screen. The monkey was trained to perform a visually guided saccade to the new (central) fixation location and to maintain gaze there for a set duration (500 ms), after which a juice reward was delivered.

During the saccade, when the eye position reached within 2.5° of the new fixation location, a probe flash was triggered. This probe, a square of size 1° × 1°, consisted of a brief luminance change (either an increase or decrease) in the pixel values of the texture at the receptive field (RF) location of the recorded superior colliculus (SC) neurons. The RFs had been previously mapped using a separate task. The receptive fields of the neurons were off the horizontal meridian, ensuring that we were measuring visual responses to peri-saccadic probes, as opposed to actual saccade-related SC motor bursts.

The flash could occur at one of three randomly chosen onset times relative to the eye crossing the 2.5° threshold: either immediately (0 ms), or with a delay of 100 ms or 200 ms. Shortly before the initial saccade onset in each trial, we calculated the fixation error and adjusted the probe position accordingly to ensure accurate alignment with the intended RF location. In total, we presented 58-75 trials per neuron per condition.

#### Data Analysis

Eye position data were recorded at 1 kHz using scleral search coils, and neuronal activity in the superior colliculus (SC) was acquired via laminar probes. Saccades were detected offline using a previously described analysis toolbox^77^, and the detected events were manually verified to ensure accuracy.

Spike sorting was performed offline using the Kilosort Toolbox^78^, followed by manual refinement with the phy software. Subsequent analyses focused on firing rates across experimental conditions. Firing rates were estimated by convolving spike trains with a Gaussian kernel (σ = 10 ms).

Only SC neurons that exhibited a significant visual response relative to baseline activity (*t*-test) were included in the analysis. This was necessary because suppression acts on visual responses. To quantify suppression strength, we employed an analysis approach similar to that used in prior work^8^.

First, to isolate the neural response associated with the saccade itself, we aligned spiking activity to the offline-detected saccade onset and averaged responses across the portion of the trial influenced by the saccade but not by the visual probe. This saccade-related response was computed separately for each neuron, saccade direction, and background texture condition.

Next, we aligned the neural activity to the onset of the probe flash and determined the temporal offset between saccade onset and probe onset for each trial. The saccade-related response was then temporally shifted by this offset and subtracted from the individual trial activity, isolating the probe-evoked component of the response.

In this study, we averaged across trials in which the probe flash was either an increase or a decrease in local luminance. Analysis was restricted to trials in which a coarse texture served as the background.

To quantify the degree of suppression, we computed a suppression index for each neuron by dividing the average evoked response in the 0 ms delay condition by the average response in the 200 ms delay condition (baseline). The evoked response was defined as the mean firing rate in the window from 50 to 200 ms after probe onset.

Finally, we note that our study quantified saccadic suppression with large saccades. However, in the SC, suppression strength does not seem to strongly depend on saccade size. For example, in Fig. 5F, our observed suppression strengths were quantitatively similar to those observed in Chen et al, 2017^16^ with much smaller microsaccades.

### Zebrafish *in-vivo* calcium imaging

#### Experimental setup

Larval zebrafish were embedded in 1.6% low-melting agarose with the agarose removed around the eyes to allow for free eye movements. The fish were then placed onto a custom-built stage and submerged in E3 within a spherical stimulation arena, previously described^10,79^. The visual stimulation arena consisted of a Ø 80 mm resin bulb filled with E3 solution with the zebrafish placed in the center. The bulb served as a projection surface for a DMD video projector, which was powered by a high-power LED light source, allowing for 360° visual stimulation. Visual stimuli were projected onto the surface of the stimulation arena at 60 Hz refresh rate at 625 nm.

To measure eye position, the fish were illuminated using a custom-built IR-LED ring and the eyes were recorded with an infrared-filtered camera (Basler ace 2 R, a2A1920-160umBAS, Basler AG, Ahrensburg, Germany) at 100 Hz. Fish were allowed to perform spontaneous saccades. Eye position was extracted online and instantaneous velocity computed. Saccades were detected, using a velocity threshold of at least 200°/s. To optimize saccade detection, the actual velocity threshold was adjusted for each fish to ensure ideal saccade detection and to avoid noise interference. To generate and display visual stimuli, to extract eye position and velocity and to trigger visual stimuli in a closed-loop manner we used *VxPy* (Version 0.1.4).

To measure calcium activity in tectal neurons we used a moveable objective two-photon setup (Sutter Instruments, Novato, CA, USA) with an acquisition rate of 2.18 fps, an image size of 512 x 512 pixels and a resolution of 0.88 μm x 0.88 μm per pixel. GCaMP6f was excited at 920 nm and emitted light was collected using a 20x/1.0 Zeiss water immersion objective.

#### Visual stimuli

Binary gaussian blur textures were displayed onto the spherical stimulation arena, as previously described^10^. To account for the spherical geometry of the zebrafish visual stimulation arena the blurry texture was generated by placing 2000 equidistant nodes onto the surface of the sphere. Half of the nodes were randomly assigned bright, the other half dark. Von-Mises-Fisher distributions with concentration, κ = 5.6 were placed centered on each bright node and contrast was rescaled to ensure roughly uniform contrast and local luminance across visual space^10^. To account for inter-species differences, this produced coarser patterns for zebrafish than for macaques, in proportion to the larger receptive fields of fish. In response to spontaneous saccades after a delay period of 20, 100, 250, 500, 1000, 2000, or 4000 ms a probe flash consisting a probe flash consisting of a whole-field luminance decrease with a temporal cosine shape. The flash started at maximal luminance and then decreased and increased luminance by 50% in a cosine profile over 500 ms (500 ms = one full cosine cycle). To obtain a baseline response to the probe flash, it was also shown 8 s after saccade detection to ensure no temporal connection between saccade and probe flash. Each peri-saccadic flash delay was shown once per trial, baseline flashes were shown twice per trial. To measure saccade-associated responses, once per trial no probe flash was shown after a saccade, yielding at total of 10 conditions per trial (7 delays, 2 baseline probe flashes, and 1 control saccade without flash). Trials were pseudo-randomized and repeated at least 4 times. Trials, in which the fish performed unwanted saccades between the initial saccade and presentation of the probe flash were discarded. Trials in which a saccade was falsely detected when the fish moved its head in a struggle motion, rather than moving its eyes, were also discarded.

#### Data Analysis

Calcium recordings were registered and segmented into ROIs using suite2p^80^. Further analysis of registered ROIs was performed with custom-written Python (PyCharm, JetBrains, Prague, Czech Republic) and Matlab (The MathWorks Inc., Natick, USA) code. Fluorescence traces of individual ROIs were normalized to ΔF/F_0_, where F_0_ was a 120 second sliding median window. ROIs that showed responses to the probe flash were identified using regression-based analysis and only those ROIs were considered for further analysis. For each ROI, saccade related responses were quantified, using control conditions, where no probe flash was shown. Said saccade-related response was subtracted in all other conditions during and shortly after the time of the saccade. Peak ΔF/F_0_ responses to probe flashes were quantified and averaged for each delay condition. For each ROI a suppression index for each delay was computed by dividing the peri-saccadic response by the baseline response, as done in macaque. For the CDF plot shown in Figure 4E, suppression indices for the delay times 20ms, 100ms and 250ms were averaged for each ROI. These delay times roughly match the saccade duration of zebrafish and thus are equivalent to the 50ms delay in macaque, which was chosen based on the saccade duration in macaques. Amongst all probe flash-responsive ROIs, truly suppressed ROIs were identified with a permutation test, described previously^10^. Briefly, data were compared to 7 potential model suppression curves, each with maximal suppression at one of the 7 different flash delays. The model suppression curve with the best correlation coefficient was confirmed through a permutation test with n = 10,000 permutations with a Bonferroni-corrected significance cut-off of p < 0.05. ROIs that reached that significance cut-off were considered “truly suppressed”.

## Acknowledgements

This work was funded by the Deutsche Forschungsgemeinschaft (DFG; German Research Foundation) through the Special Priority Programme: “SPP 2205 Evolutionary optimization of neural processing” (430158665 and 430157666; AR 1076/1-2, HA 6749/3-2, AR 1076/1-1), by the Major Research Instrumentation programme of the DFG (INST 37/1254-1 FUGG), and by the Human Frontier Science Program (HFSP) Young Investigator grant RGY0079. We are grateful to Frédéric Crevecoeur of UC Louvain, Belgium, for helpful discussions of the original Bayesian estimator model, as well as for sharing some of its unpublished code.

## Author Contributions

G.S. performed experiments in zebrafish optic tectum, analyzed zebrafish calcium data, generated figures, and wrote the manuscript and all major revisions. F.A.D. reproduced the saccadic suppression model, generated figures, and wrote the manuscript. M.P.B. performed macaque experiments, analyzed macaque eye movement data, analyzed macaque electrophysiology data, generated figures, and wrote the manuscript. I.T. analyzed zebrafish and macaque eye trace data and generated figures. Y.Y. performed macaque experiments. T.M. performed macaque experiments and analyzed macaque electrophysiology data. Z.M.H. and A.B.A. designed the overall study, acquired funding and wrote the manuscript.

## Resource availability

### Lead contacts

Aristides B. Arrenberg, Ziad M. Hafed.

### Materials availability

Materials are available from the authors upon request.

### Data and code availability

Zebrafish data and part of the code are available at in a public repository (https://doi.gin.g-node.org/…). Macaque code and data as well as the code for the model are available on Github (https://github.com/…).

